# Improved molecular characterization of the *Klebsiella oxytoca* complex reveals the prevalence of the kleboxymycin biosynthetic gene cluster

**DOI:** 10.1101/2020.07.24.215400

**Authors:** Preetha Shibu, Frazer McCuaig, Anne L. McCartney, Magdalena Kujawska, Lindsay J. Hall, Lesley Hoyles

## Abstract

As part of ongoing studies with clinically relevant *Klebsiella* spp., we characterized the genomes of three clinical GES-5-positive ST138 strains originally identified as *Klebsiella oxytoca*. *bla*_OXY_ gene, average nucleotide identity and phylogenetic analyses showed the strains to be *Klebsiella michiganensis*. Affiliation of the strains to ST138 led us to demonstrate that the current multi-locus sequence typing scheme for *K. oxytoca* can be used to speciate members of this genetically diverse complex of bacteria. The strains encoded the kleboxymycin biosynthetic gene cluster (BGC), previously only found in *K. oxytoca* strains and one strain of *Klebsiella grimontii*. The finding of this BGC, associated with antibiotic-associated haemorrhagic colitis, in *K. michiganensis* led us to carry out a wide-ranging study to determine the prevalence of this BGC in *Klebsiella* spp. Of 7,170 publicly available *Klebsiella* genome sequences screened, 88 encoded the kleboxymycin BGC. All BGC-positive strains belonged to the *K. oxytoca* complex, with strains of four (*K. oxytoca*, *K. pasteurii*, *K. grimontii*, *K. michiganensis*) of the six species of the complex found to encode the complete BGC. In addition to being found in *K. grimontii* strains isolated from preterm infants, the BGC was found in *K. oxytoca* and *K. michiganensis* metagenome-assembled genomes recovered from neonates. Detection of the kleboxymycin BGC across the *K. oxytoca* complex may be of clinical relevance and this cluster should be included in databases characterizing virulence factors, in addition to those characterizing BGCs.

**IMPACT STATEMENT:** Members of the *Klebsiella oxytoca* complex are difficult to speciate using phenotypic and chemotaxonomic methods. Consequently, many genomes deposited in public databases are misclassified as *K. oxytoca*. Here we demonstrate that the current multi-locus sequence typing (MLST) system for the complex can be used to accurately speciate many strains, which will be of use to clinical laboratories in resource-limited settings which rely on the MLST scheme for typing and epidemiological tracking of isolates. In addition, extended analyses of the genomes of *Klebsiella* spp. have revealed the kleboxymycin biosynthetic gene cluster (BGC) is restricted to species of the *Klebsiella oxytoca* complex (*K. oxytoca*, *K. michiganensis*, *K. pasteurii* and *K. grimontii*). Species- and/or gene-specific differences in the cluster’s sequences may be relevant to virulence of *K. oxytoca* and related species. The finding of the kleboxymycin BGC in the preterm infant gut microbiota may have implications for disease presentation in a subset of neonates.

## INTRODUCTION

Members of the *Klebsiella oxytoca* complex encode a chromosomal β-lactamase gene (*bla*_OXY_) (1). Differences in the sequence of this gene allowed the establishment of phylogroups (Ko), which correspond to species: *K. michiganensis* (Ko1, with Ko5 representing a sub-lineage), *K. oxytoca* (Ko2), *K. spallanzanii* (Ko3), *K. pasteurii* (Ko4), *K. grimontii* (Ko6) and *K. huaxiensis* (Ko8). Ko7 has been described on the basis of a single isolate (1). Individual gene (*rpoB*, *gyrA*, *rrs*) sequences can be used to differentiate species of the complex (2), as can genome-based average nucleotide identity (ANI) and phylogenomic analyses (1,3). All members of the *K. oxytoca* complex can be differentiated by MALDI-TOF (1), but reference databases currently in routine clinical use lack reference spectra of the different species to allow identification beyond *K. oxytoca*.

Recent work has demonstrated genomic characterization of *K. oxytoca* strains is inadequate, with large numbers of genomes deposited in public databases erroneously assigned to *K. oxytoca* instead of *K. michiganensis* or *K. grimontii* (3–6). Consequently, *K. michiganensis* and *K. grimontii* are clinically relevant but under-reported in the literature (3,7). Given that the *bla*_OXY_ gene has diversified in parallel to housekeeping genes in the *K. oxytoca* complex, it is likely that the *K. oxytoca* multi-locus sequence typing (MLST) scheme (8) can be used to speciate all members of this genetically diverse group of bacteria.

Little is known about the antibiotic-resistance and virulence genes encoded by *K. oxytoca* and related species. In the course of ongoing *Klebsiella*–phage work, with three GES-5-positive ST138 strains originally described as *K. oxytoca* (9,10), we sought to determine whether widely recognized virulence factors such as enterobactin, yersiniabactin and salmochelin are encoded in the strains’ genomes, and the kleboxymycin biosynthetic gene cluster (BGC), as this was until recently a little-studied BGC implicated in non-*Clostridioides difficile* antibiotic-associated haemorrhagic colitis (AAHC) (11–14). AAHC is caused by the overgrowth of cytotoxin-producing *K. oxytoca* secondary to use of antibiotics such as penicillin or amoxicillin, resulting in the presence of diffuse mucosal oedema and haemorrhagic erosions (15,16). This type of colitis is distinct from the more common form of antibiotic-associated diarrhoea caused by toxin-producing *Clostridiodes difficile*, which usually gives rise to watery diarrhoea resulting in mild to moderate disease.

Gene-based and genomic analyses of our ST138 isolates showed they were *K. michiganensis*, not *K. oxytoca*, and that along with common virulence genes they encoded the kleboxymycin BGC. Our findings led us to 1) determine whether the *K. oxytoca* MLST scheme could be used to speciate members of the *K. oxytoca* complex, and 2) investigate the distribution of the kleboxymycin BGC in a range of *Klebsiella* and related species.

## METHODS

### Clinical isolates

Strains PS_Koxy1 (isolated December 2014; cardiothoracic/intensive care unit), PS_Koxy2 (isolated August 2015; haematology unit) and PS_Koxy4 (isolated September 2015; haematology unit) had been recovered from a throat swab, urine and rectal swab, respectively, obtained from three different adults. The strains were from the study of Eades *et al.* (9), described in further detail by Ellington *et al.* (10) (PS_Koxy1, patient X; PS_Koxy2, patient A; PS_Koxy4, patient B; Frances Davies, personal communication). The study of anonymized isolates beyond the diagnostic requirement was approved by an NHS research ethics committee (number 06/Q0406/20). Full details of Methods associated with the phenotypic and genotypic characterization of the clinical isolates can be found in **Supplementary Material**.

### ANI analysis of genome sequences

All annotated non-redundant *Klebsiella* genome assemblies available in the NCBI Genome database on 2 September 2019 (*n* = 7,170; **Supplementary Table 1**) were downloaded (17). ANI of genomes with their closest relatives and type strains of species was assessed using FastANI (18), which uses Mashmap as its MinHash-based alignment-free sequence mapping engine to provide ANI values for both complete and draft-quality genomes that are related by 80–100 % ANI.

### MLST analyses

Allele sequences (<u>*n*=442</u> representing seven housekeeping genes – *gapA*, *infB*, *mdh*, *pgi*, *phoE*, *rpoB*, *tonB* – contributing to 354 different MLST sequence types; correct as of 19 March 2021) for the *K. oxytoca* MLST scheme (8) were used to determine the MLST profiles of all *K. oxytoca* complex genomes included in this study (**Supplementary Table 1**). The allele sequences were used to create BLASTN databases against which the assemblies of all genomes included in this study were searched. Sequences with exact hits to one allele of each housekeeping gene were retained, allowing us to identify the sequence types of the genomes included in this study (**Supplementary Table 2**). For those genomes that returned hits to alleles across all seven housekeeping genes, a phylogenetic tree (neighbour joining, Jukes Cantor) was generated in Geneious Prime v2019.2.1 using the aligned (CLUSTAL W) concatenated (*gapA*–*infB*–*mdh*–*pgi*–*phoE*–*rpoB*–*tonB*) nucleotide sequences of their housekeeping genes and those of each sequence type used in the *K. oxytoca* MLST scheme (8). Support for clustering of nodes in the tree was determined by bootstrap analysis (1,000 replications).

### Characterization of the kleboxymycin BGC in genomes

The annotated reference sequence of the kleboxymycin BGC was downloaded from GenBank (accession number MF401554 (11)) and used as a BLASTP database for searches with the protein sequences encoded within the genomes of PS_Koxy1, PS_Koxy2 and PS_Koxy4. Initially, Geneious Prime v2019.2.1 was used to identify regions of the three draft genomes encoding the complete BGC, and to align them to MF401554.

The protein sequences of the annotated assemblies were searched for the kleboxymycin BGC using the reference sequence and BLASTP v2.9.0+, and the resulting hits were filtered based on >70 % identity and >70 % coverage to identify isolates potentially carrying genes from the BGC. *K. grimontii* (*n*=3) and *K. michiganensis* (*n*=2) and *K. oxytoca*-related metagenome-assembled genomes (MAGs) (*n*=25) from Chen *et al.* (3) were also subject to BLASTP searches. Genomes that encoded the full BGC (i.e. all 12 BGC genes on a contiguous stretch of DNA) were identified from the BLAST results. The protein sequences encoded in the BGC were extracted from the annotated assemblies using samtools v1.9 faidx (19) and concatenated into a single sequence (the sequence data are available as supplementary material from <u>figshare</u>). These concatenated sequences were used to produce a multiple-sequence alignment (MSA) in Clustal Omega v1.2.4, along with the BGC sequences of the three *K. michiganensis* clinical isolates, the reference sequence (11), a recently described *K. grimontii* sequence (20) and a homologous sequence found in *Pectobacterium brasiliense* BZA12 (to be used as an outgroup in later phylogenetic analyses; identified as encoding the complete kleboxymycin BGC through NCBI BLASTP). Phylogenetic analyses were carried out on the MSA using the R package Phangorn v2.5.5 (21), producing a maximum-likelihood tree, which was visualised and rooted (on *P. brasiliense* BZA12) using the Interactive Tree of Life (iTOL v5.5) (22). To examine variation at the individual protein level, further within-species MSAs were produced for each of the 12 protein sequences in the BGC. Each of these alignments was used as the basis for a consensus sequence, produced using EMBOSS Cons v6.6.0.0, representing each of the four species carrying the BGC. An MSA and per cent identity matrix were then generated for each protein between the consensus sequences of *K. oxytoca*, *K. grimontii*, *K. michiganensis* and *K. pasteurii*, along with the reference sequence (11).

The species affiliations of the genomes encoding the full kleboxymycin BGC were determined using FastANI v1.2 (18) against genomes of type strains of the *K. oxytoca* and *K. pneumoniae* complexes (1,23) and *K. aerogenes* ATCC 13048^T^ (assembly accession number GCA_003417445), with PhyloPhlAn 0.99 used to conduct a phylogenetic analysis to confirm species affiliations. PhyloPhlAn identifies hundreds of conserved (core) proteins from a given genomic dataset and uses them to build a complete high-resolution phylogeny.

## RESULTS

### Characterization of the clinical isolates

Although initial phenotypic tests (**Supplementary Material**) and genomic analyses (9,10) identified PS_Koxy1, PS_Koxy2 and PS_Koxy4 as *K. oxytoca*, analyses of the isolates’ proteomes showed them to be *K. michiganensis* ST138 (phylogroup Ko1, *bla*_OXY1-8_) (**Supplementary Figure 1**). Full details of phenotypic characterization and genome sequencing of the clinical isolates can be found in **Supplementary Material**. PS_Koxy1, PS_Koxy2 and PS_Koxy4 all shared 98.81 %, 98.71 % and 98.71 % ANI, respectively, with the type strain of *K. michiganensis* (W14^T^, GCA_901556995), and 99.98 to 100.00 % ANI with each other. Based on current recommendations, ANI of 95–96 % and above with the genome of the type strain is indicative of species affiliation (24). Inclusion of the genomes with representatives of all six species of the *K. oxytoca* complex in a phylogenetic analysis confirmed the affiliation of PS_Koxy1, PS_Koxy2 and PS_Koxy4 with *K. michiganensis* (**Supplementary Figure 2**).

### Assigning MLST sequence types to species

While annotations for genomes are improving, we have previously noted and continue to notice issues with identities attributed to *K. oxytoca* genomes in public repositories (3). Consequently, the identity of all genomes included in this work was first confirmed by ANI analysis (**Supplementary Table 1**), with *bla*_OXY_ gene and phylogenetic analyses supporting our findings (**Supplementary Material**). Of the 178 *K. oxytoca* complex genomes identified, many had been misassigned in GenBank: seven genomes were listed as *K. grimontii*, 106 as *K. oxytoca*, 51 as *K. michiganensis*, 13 as *Klebsiella* sp. and one as *K. pneumoniae*. Our analyses of the 178 genomes showed the dataset actually represented *K. michiganensis* (*n*=76), *K. oxytoca* (*n*=66), *K. grimontii* (*n*=24), *K. pasteurii* (n=6), *K. huaxiensis* (*n*=5) and *K. spallanzanii* (*n*=1).

The *K. oxytoca* MLST scheme uses sequence polymorphisms among seven housekeeping genes – *gapA*, *infB*, *mdh*, *pgi*, *phoE*, *rpoB*, *tonB* – to generate sequence types for isolates. Currently, there are 442 allele sequences that contribute to 354 unique MLST sequence types. We first identified nucleotide sequences within the genomes with exact matches to nucleotide sequences within the allele reference dataset. One-hundred-and-twenty-nine genomes returned hits to known MLST profiles, and 10 isolates returned MLST profiles with no assigned sequence type (**Supplementary Table 2**). Our clinical isolates returned the expected ST138 result.

Of the 66 *K. oxytoca* genomes, 59 could be assigned to known sequence types (in order of abundance: ST2, ST176, ST199, ST36, ST19, ST30, ST53, ST101, ST18, ST31, ST34, ST48, ST58, ST59, ST141, ST145, ST153, ST181, ST221, ST222, ST257, ST258, ST287, ST323) and one (GCA_003937225) represented a novel sequence type. Of the 24 *K. grimontii* genomes, 13 could be assigned to known sequence types (ST172, ST216, ST104, ST186, ST236, ST263, ST316, ST319, ST350), with four (GCA_002856195, GCA_900451335, GCA_008120915, GCA_004343645) representing unique novel sequence types. Of the six *K. pasteurii* genomes, three could be assigned to known sequence types (ST47, ST311, ST351) and one (GCA_901563825) represented a novel sequence type. Of the 79 *K. michiganensis* genomes (including our three clinical isolates), 57 could be assigned to known sequence types (ST85, ST27, ST202, ST143, ST29, ST50, ST84, ST138, ST11, ST88, ST317, ST28, ST40, ST52, ST82, ST92, ST98, ST108, ST127, ST144, ST146, ST157, ST170, ST180, ST226, ST294, ST315), with four genomes (GCA_000783895, GCA_000735215, GCA_007097185, GCA_007097115) representing three novel sequence types. None of the *K. huaxiensis* or *K. spallanzanii* genomes returned hits to known alleles (**Supplementary Table 2**), but the relevant individual housekeeping gene sequences are provided as **Supplementary Files** for use by other researchers.

For those genomes that encoded known or novel sequence types, we concatenated their housekeeping-gene sequences and used them to create a MSA with the concatenated sequences of each of the 354 recognized MLST sequence types. This MSA was used to create a phylogenetic tree, allowing us to visualize the relationships among species and sequence types (**Figure 1**).

**Figure 1.**
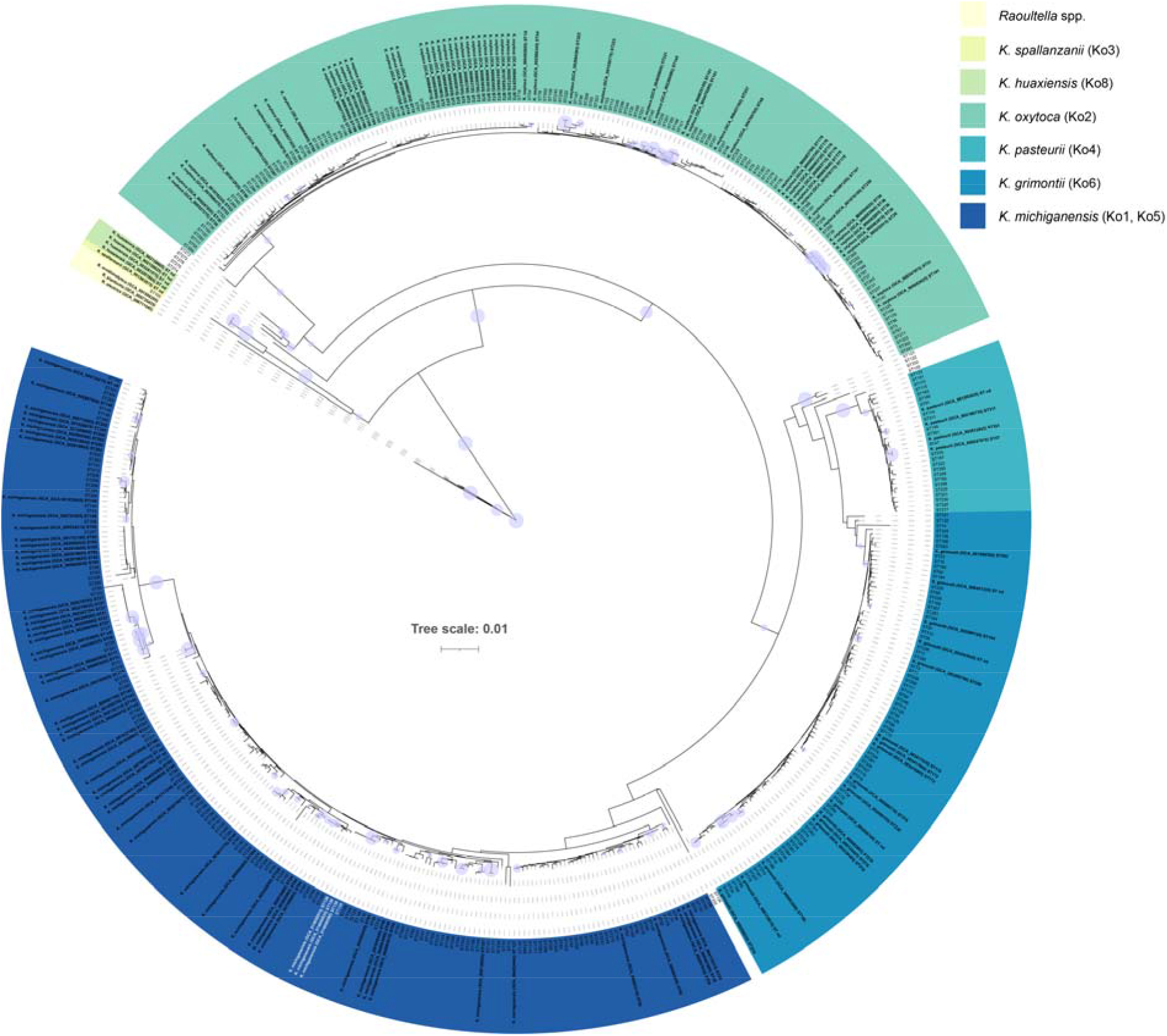
Sequence types within the *K. oxytoca* MLST scheme can be used to speciate members of the *K. oxytoca* complex. The three *K. michiganensis* clinical isolates (all ST138) characterized in this study are shown in white. The phylogenetic tree (neighbour joining, Jukes Cantor) was generated using concatenated nucleotide sequences of housekeeping genes (*gapA*–*infB*–*mdh*–*pgi*–*phoE*–*rpoB*– *tonB*) used in the *K. oxytoca* MLST scheme (8). The purple circles represent bootstrap values ≥80 % (based on 1,000 replications); the larger the circle, the higher the bootstrap value. Scale bar, average number of nucleotide substitutions per position. The full list of MLST sequence types and their species affiliations are available in **Supplementary Table 2**.

Of the 354 known MLST sequence types, 342 (96.6 %) were associated with specific members of the *K. oxytoca* complex (**Supplementary Table 2**): 115 with *K. oxytoca*, 130 with *K. michiganensis*, 73 with *K. grimontii* and 24 with *K. pasteurii*. Eleven were associated with unspecified members of the *K. oxytoca* complex. ST105 was associated with *Raoultella ornithinolytica*, sharing 99.73 % sequence similarity type strain’s MLST profile. *K. oxytoca*-specific sequence types shared 98.64–100 % sequence similarity, *K. michiganensis*-specific sequence types shared 96.62–100.00 % sequence similarity, *K. grimontii*-specific sequence types shared 98.20–100.00 % sequence similarity, *K. pasteurii*-specific sequence types shared 99.00–100.00 % sequence similarity and *K. huaxiensis*-specific sequence types shared 97.09–99.7 % sequence similarity. A matrix of similarity values for the 504 sequences included in the analysis is available in **Supplementary Material**, along with the MSA alignment used to generate the phylogenetic tree shown in **Figure 1**.

### Detection of the complete kleboxymycin BGC in clinical isolates

It has long been known that *K. oxytoca* gut colonization is linked with AAHC (16). Schneditz *et al.* (12) showed tillivaline (TV), a pyrrolobenzodiazepine (PBD) derivative produced by *K. oxytoca*, is one of the enterotoxins responsible for causing AAHC. This toxic product is encoded by the heterologous expression of the kleboxymycin [also known as tilimycin (TM) (14)] BGC comprising 12 genes (11). Protein sequences of the reference sequence (11) were used to create a BLASTP database against which the proteins encoded in the genomes of PS_Koxy1, PS_Koxy2 and PS_Koxy4 were compared. The genomes of PS_Koxy1, PS_Koxy2 and PS_Koxy4 encoded a complete kleboxymycin BGC (**Figure 2**). All genes in each of the genomes shared >99 % identity and >99 % query coverage with the genes of the reference sequence (12): *mfsX*, 99.76 % identity; *uvrX*, 99.87 %; *hmoX*, 99.80 %; *adsX*, 99.85 %; *icmX*, 99.52 %; *dhbX*, 99.62 %; *aroX*, 99.74 %; *npsA*, 99.80 %; *thdA*, 98.68 %; *npsB*, 99.93 %; *npsC*, 98.47 %; *marR*, 99.39.

**Figure 2.**
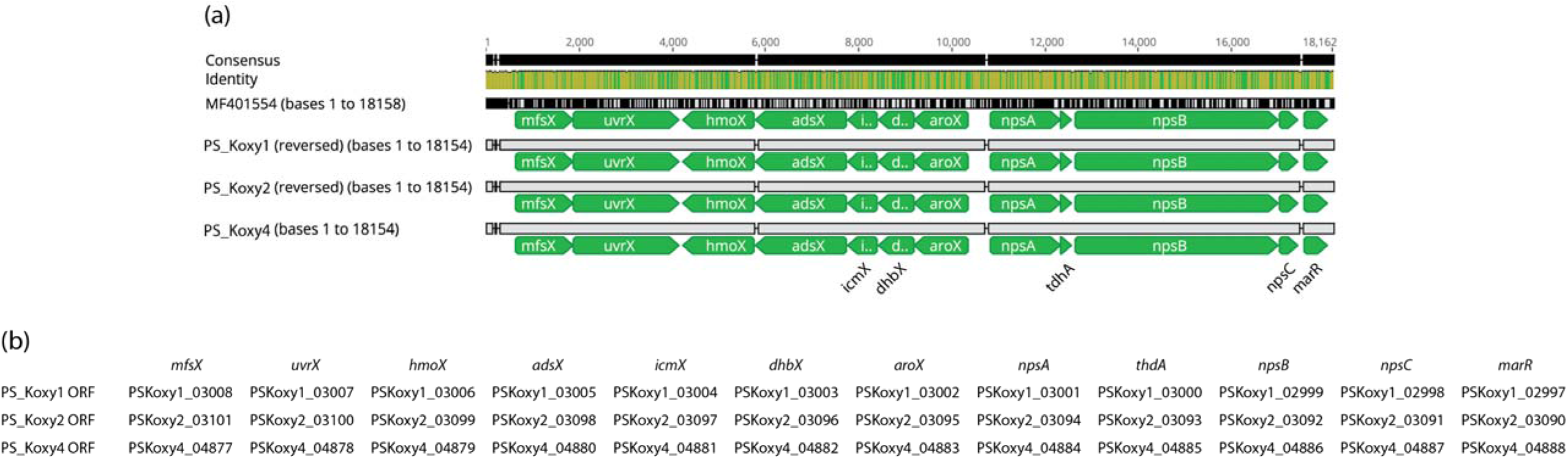
Alignment of the kleboxymycin BGCs from the three clinical *K. michiganensis* strains with the complete cluster of *K. oxytoca* MH43-1 [GenBank accession number MF401554 (11)]. (a) The image (alignment view) was generated via the progressiveMauve algorithm plugin of Geneious Prime v2019.2.1 (default settings, full alignment), with gene names for the three clinical isolates assigned manually. (b) Genes corresponding to Prokka-generated annotations. Consensus identity is the mean pairwise nucleotide identity over all pairs in the column: green, 100 % identity; greeny-brown, at least 30 % and under 100 % identity; red, below 30 % identity.

Our strains were *K. michiganensis* ST138, so we downloaded and assembled (from BioProject <u>PRJEB30858</u>) available raw sequence data from 19 *K. oxytoca* ST138 strains described recently (10) and determined whether they were in fact *K. michiganensis* and encoded the kleboxymycin BGC. All strains were confirmed to be *K. michiganensis* on the basis of ANI analysis, and encoded the complete kleboxymycin BGC (**Supplementary Figure 4**).

Schneditz *et al.* (12) reported *npsA*/*npsB* were functionally conserved in six sequenced strains of *K. oxytoca* (**Table 1**), based on a BLASTP analysis. Full details of the analysis are unavailable, with only a brief mention of presence being determined based on BLASTP sequence identities >90 % with no indication of sequence coverage. All the genomes included in the study of Schneditz *et al.* (12) were compared with those of the type strain of *K. oxytoca* and related species to confirm their species affiliations (**Table 1**). While some strains were *K. oxytoca*, others belonged to *K. michiganensis*, *K. pasteurii*, *K. grimontii* and *R. ornithinolytica*. Using thresholds of 70 % identity and 70 % query coverage in our BLASTP analyses to reduce the potential for detecting false positives, we reanalysed the genomes included in the study of Schneditz *et al.* (12). Our results agreed with those of Schneditz *et al.* (12) for all genomes, except we detected *npsA*/*npsB* (and all other genes encoded in the kleboxymycin BGC) in *K. grimontii* SA2. *K. oxytoca* 10–5243, *K. pasteurii* 10–5250, *K. oxytoca* 11492-1, *K. oxytoca* 10–5248 and *K. grimontii* M5a1 also encoded the whole kleboxymycin BGC. All genes in all matches shared greater than 90 % identity across greater than 99 % query coverage. *K. michiganensis* 10–5242, E718 and KCTC 1686 did not encode homologues associated with the kleboxymycin BGC. *K. oxytoca* 10–5245 encoded almost-complete homologues of four genes [EHS96696.1 (*marA*) 98.79 % identity, 99.39 % coverage; EHS96697.1 (*npsC*) 95.38 % identity, 99.23 % coverage; (EHS96698.1 (*mfsX*) 96.68 % identity, 99.87 % coverage; EHS96699.1 (*uvrX*) 94.88 % identity, 99.76 % coverage] in contig JH603137.1.

**Table 1.** Genomes included in analyses conducted by Schneditz *et al.* (12) with corrected species affiliations (originally reported as *K. oxytoca*)

| Assembly accession | Strain | Species | ANI with shown genome* | *npsA/npsB* |
| --- | --- | --- | --- | --- |
| GCA_000240325.1 | KCTC 1686 | *K. michiganensis* | 98.69 %, GCA_901556995.1 | – |
| GCA_000247835.1 | 10–5242 | *K. michiganensis* | 97.59 %, GCA_901556995.1 | – |
| GCA_000247855.1 | 10–5243 | *K. oxytoca* | 99.31 %, GCA_900977765.1 | + |
| GCA_000247875.1 | 10–5245 | *K. oxytoca* | 99.13 %, GCA_900977765.1 | – |
| GCA_000247895.1 | 10–5246 | *Raoultella ornithinolytica* | 99.21 %, GCA_001598295.1 | – |
| GCA_000247915.1 | 10–5250 | *K. pasteurii* | 99.29 %, GCA_901563825.1 | + |
| GCA_000252915.3 | 11492-1 | *K. oxytoca* | 99.15 %, GCA_900977765.1 | + |
| GCA_000276705.2 | E718 | *K. michiganensis* | 98.37 %, GCA_901556995.1 | – |
| GCA_000427015.1 | SA2 | *K. grimontii* | 99.33 %, GCA_900200035.1 | + |
| GCA_001078235.1 | 10–5248 | *K. oxytoca* | 99.25 %, GCA_900977765.1 | + |
| GCA_001633115.1 | M5a1 | *K. grimontii* | 99.40 %, GCA_900200035.1 | + |

\*GCA\_901556995.1 = *K. michiganensis* W14<sup>T</sup>; GCA\_900200035.1, *K. grimontii* 06D021<sup>T</sup>; GCA\_900977765.1 = *K. oxytoca* ATCC 13182<sup>T</sup>; GCA\_001598295.1 = *R. ornithinolytica* NBRC 105727<sup>T</sup>; *K. pasteurii* SB3355<sup>T</sup> GCA\_901563825.1.

### Detection of the kleboxymycin BGC in the faecal microbiota of preterm infants

Our previous work had highlighted the preterm infant gut microbiota harbours a range of species belonging to the *K. oxytoca* complex (3). BLASTP searches of the two *K. michiganensis* (P049A W, GCA_008120305; P095L Y, GCA_008120085) and three *K. grimontii* (P038I, GCA_008120465; P043G P, GCA_008120425; P079F P, GCA_008120915) strains we previously characterized showed all three *K. grimontii* strains encoded the kleboxymycin BGC (**Supplementary Figure 5**). All BGC genes in their genomes shared >98 % identity and >99 % query coverage with the genes of the reference sequence (11): *mfsX*, 100 % identity; *uvrX*, 99.60–99.73 %; *hmoX*, 99.80 %; *adsX*, 99.69 %; *icmX*, 100 %; *dhbX*, 100 %; *aroX*, 99.94–99.74 %; *npsA*, 99.21–99.41 %; *thdA*, 98.68 %; *npsB*, 99.31–99.38 %; *npsC*, 99.23–100 %; *marR*, 100 %. The BGC was also detected in 8/25 of the preterm-associated *K. oxytoca* complex MAGs (3 *K. oxytoca*, 5 *K. michiganensis*) we described previously (3). An MSA of the preterm-associated genomes’ BGC against the reference sequence (11) suggested species-specific clustering of the sequences (**Supplementary Figure 5**).

### Prevalence of the kleboxymycin BGC in *Klebsiella* spp

Given the work detailed above had detected the kleboxymycin BGC in several different but closely related *Klebsiella* species and in a range of clinical and gut-associated isolates, and Hubbard *et al.* (20) recently detected the BGC in a strain of *K. grimontii*, we chose to increase the scope of our analysis to include 7,170 publicly available assembled *Klebsiella* genomes (including our three clinical strains, and five isolates from preterm infants (3)) (**Supplementary Table 1**).

As mentioned above, we have noted issues with identities attributed to *Klebsiella* genomes in public repositories (3), so the identity of all non-*K. oxytoca* complex genomes included in this work was first confirmed by ANI analysis (**Supplementary Table 1**). The majority (*n*=6,245) of the additional genomes were *K. pneumoniae*, followed by *K. variicola* subsp. *variicola* (*n*=241), *K. quasipneumoniae* subsp. *similipneumoniae* (*n*=184), *K. aerogenes* (*n*=168), *K. quasipneumoniae* subsp. *quasipneumoniae* (*n*=120), *K. variicola* subsp. *tropica* (*n*=19), ‘*K. quasivariicola*’ (*n*=11) and *K. africana* (*n*=1). Out of 7,170 genomes, 110 (1.5 %) had one or more matches with the 12 genes encoded within the kleboxymycin BGC reference sequence, with all except two genomes (both *K. pneumoniae*) belonging to species of the *K. oxytoca* complex (**Supplementary Table 3**). Ninety-six genomes – all belonging to the *K. oxytoca* complex – encoded at least 12 genes belonging to the BGC (**Supplementary Table 3**), and were examined further.

One genome (GCA_002856195) encoding 12 BGC genes was found to encode two stretches of the same protein with the other cluster-associated genes non-contiguous, while one (GCA_004005605) encoded 13 BGC genes (one gene duplicated) in a non-contiguous arrangement. Fifty-five out of 66 (83.3 %) *K. oxytoca* genomes encoded the entire kleboxymycin BGC, as did 19/24 (79.2 %) *K. grimontii*, 9/79 (11.4 %) *K. michiganensis* and 5/6 (83.3 %) *K. pasteurii* genomes (**Figure 3a**). Phylogenetic analysis (**Figure 3b**) confirmed findings from ANI analyses (**Supplementary Table 1**) that showed all genomes belonged to species of the *K. oxytoca* complex. The 88 genomes confirmed to encode the complete kleboxymycin BGC included the type strain of *K. grimontii*. The BGC cluster sequences grouped according to species, and the reference sequence (11) clustered with *K. grimontii* sequences and was closely related to the type strain of that species (**Figure 3c**).

**Figure 3.**
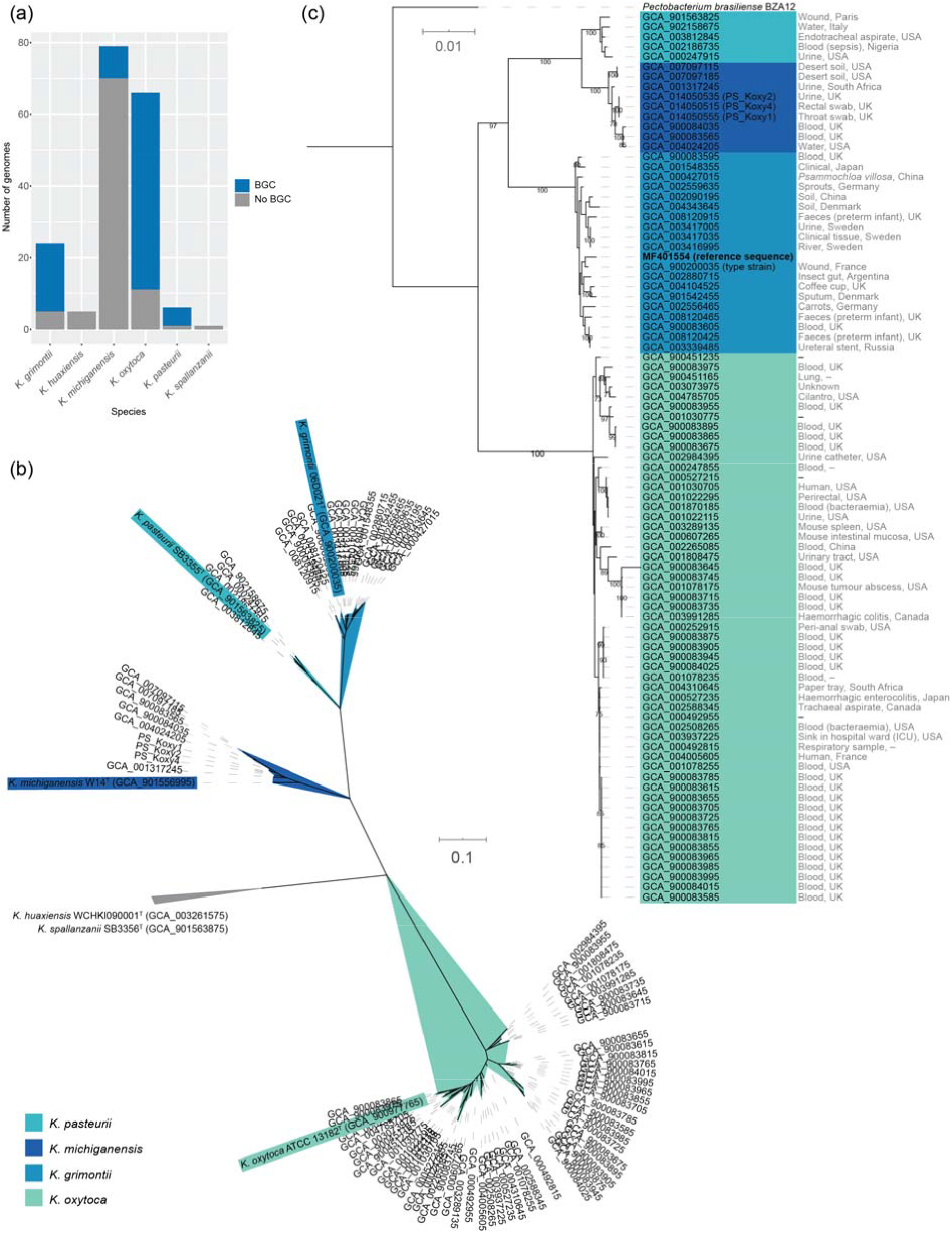
Distribution of the kleboxymycin BGC in *Klebsiella* spp. genomes. (a) Distribution of the *K. oxytoca* complex genomes encoding the entire kleboxymycin BGC. (b) Unrooted maximum-likelihood tree [generated using PhyloPhlAn v0.99 (34) and 380 protein-encoding sequences conserved across the genomes] confirming species affiliations of the 88 genomes within the *K. oxytoca* complex (1) encoding the kleboxymycin BGC. Type strains are shown with coloured backgrounds corresponding to the legend. The clade associated with *K. huaxiensis* and *K. spallanzanii* has been collapsed because of space constraints. (c) Maximum-likelihood tree generated with the concatenated protein sequences for the kleboxymycin BGC of the 88 genomes found to encode all 12 genes of the BGC plus the reference sequence (11). The tree was rooted using the kleboxymycin-encoding BGC of *Pectobacterium brasiliense* BZA12. Values at nodes, bootstrap values expressed as a percentage of 100 replicates. Sources of isolates, where known, are shown to the right of the assembly accession numbers. (b, c) Scale bar, average number of substitutions per position.

Species-specific consensus sequences were generated for all genes within the kleboxymycin BGC and are available as supplementary material from <u>figshare</u>. Similarity values for each gene within the BGC consensus sequences across the four species are available in **Supplementary Table 4**.

## DISCUSSION

### Genotypic characteristics of the three clinical *K. michiganensis* strains

The three clinical strains characterized herein had previously been included in a study of outbreak strains encoding GES-5 and CTX-M-15 (9), the first report of GES-5-positive clinical isolates of *K. oxytoca* ST138 in the UK. Subsequently, it has been shown that the GES-5 gene in these strains is encoded on an IncQ group plasmid (10). The whole-genome sequence data reported on previously (9) were not available to us. API 20E (this study; **Supplementary Material**), MALDI-TOF and limited sequence analysis (9) had shown the strains to be *K. oxytoca*. Our previous work with isolates recovered from preterm infants had shown that API 20E testing on its own was insufficient to accurately identify *K. oxytoca* strains (3). The strains described by Eades *et al.* (9) were characterized before the availability of MALDI-TOF databases capable of splitting species of the *K. oxytoca* complex (MALDI-TOF was only able to identify as *K. oxytoca* but did not have sufficient resolution to identify individual species within the complex) (1). As we are using PS_Koxy1, PS_Koxy2, PS_Koxy4 in ongoing phage work, we generated draft genome sequences for the strains, to accurately identify them and facilitate detailed host–phage studies in the future.

ANI and phylogenetic analyses confirmed all three strains belonged to the species *K. michiganensis*, not *K. oxytoca* (**Supplementary Material**). In addition to the AMR genes GES-5 (β-lactamase with carbapenemase activity) and CTX-M-15 (an ESBL responsible for resistance to cephalosporins) reported previously (9), the strains encoded SHV-66, an ESBL not previously reported in *K. oxytoca* and related species (**Supplementary Material**). SHV-66 has previously only been reported in a minority of β-lactamase-producing *K. pneumoniae* in Guangzhou, China (25). In this study, SHV-66 (99.65 % identity, bit-score 580 – strict CARD match) was also found in *K. michiganensis* strains E718 (26), GY84G39 (unpublished), K1439 (unpublished) and 2880STDY5682598 (7) (accession numbers GCA_000276705, GCA_001038305, GCA_002265195 and GCA_900083915, respectively), included in the phylogenetic analysis shown in **Supplementary Figure 1**. Moradigaravand *et al.* (7) noted in their study that 2880STDY5682598 encoded a *bla*_SHV_ gene, but did not document its type nor indicate its novelty.

The three strains had identical virulence factor profiles (**Supplementary Figure 3b**), encoding the plasminogen activating omptin Pla, the Mg^2+^ transport proteins MgtBC, Hsp60, autoinducer-2 (LuxS), type I fimbriae, type 3 fimbriae, type 6 secretion system I, *Escherichia coli* common pilus and enterobactin. They also encoded numerous proteins associated with capsule, regulation of capsule synthesis (RcsAB) and LPS, with several of the latter sharing identity with *Haemophilus* endotoxins (RfaD, GalU, LpxC, GmhA/LpcA, KdsA). All six proteins required for allantoin utilization were encoded in the strains’ genomes.

No capsule or O antigen types could be assigned to the strains using Kaptive, but all three strains were best matched with KL68 [PS_Koxy1, 17/18 genes matched (*cpsACP* missing); PS_Koxy2 and PS_Koxy4, 16/18 genes matched (*cpsACP* and KL68_18 missing)] and O1v1 [4/7 genes (*wzm*, *wzt*, *glf*, *wbbO*) matched in all strains].

### MLST sequence types can be used to speciate members of the *K. oxytoca* complex

The *bla*_OXY_ gene diversified in parallel to housekeeping genes in the *K. oxytoca* complex, and it is already known that *rpoB* – one of the seven genes included in the *K. oxytoca* MLST scheme (8) – can be used to speciate members of the complex (2). Given that our three clinical strains were ST138 and belonged to *K. michiganensis*, we determined whether specific sequence types within the MLST scheme could be assigned to species. We found that all species of the *K. oxytoca* complex are associated with specific sequence types. In addition, we identified 10 novel MLST sequence types that can be used to identify *K. grimontii*, *K. michiganensis*, *K. oxytoca* and *K. grimontii* genomes (**Supplementary Table 2**).

Herzog *et al.* (27), when originally describing the *K. oxytoca* MLST scheme to characterize clinical isolates, showed their concatenated sequence data for 74 clinical *K. oxytoca* isolates were associated with three clusters (A, B1 and B2). Comparison of their sequence types with our annotations shows that cluster A represents *K. oxytoca*, cluster B1 represents *K. michiganensis* and cluster B2 represents *K. grimontii* and *K. pasteurii*.

The ability to use the *K. oxytoca* MLST scheme to speciate clinical isolates will be of particular interest to clinical microbiologists in resource-limited settings who rely on the MLST scheme for typing and epidemiological tracking of isolates in the absence of whole-genome sequence data. It should also be noted that ribosomal MLST (28) (rMLST) available via the <u>Species ID portal</u> of the PubMLST website allows those working with genome sequence data derived from *K. oxytoca* complex isolates to speciate isolates. This resource uses 53 genes encoding the bacterial ribosome protein subunits (*rps* genes) to rapidly characterize genomic data to the species level.

The identification of ST105 as belonging to *Raoultella ornithinolytica* indicates this sequence type should be withdrawn from the *K. oxytoca* MLST scheme.

### Distribution of the kleboxymycin BGC in *Klebsiella* spp

As relatively little is known about the virulence factors of *K. oxytoca* and related species, and the VFDB is limited with respect to the number of *Klebsiella* spp. on which it reports information, we wanted to see whether our strains encoded the kleboxymycin BGC responsible for generating microbiome-associated metabolites known to directly contribute to AAHC (11,12). The cytotoxic nature of a heat-stable, non-proteinaceous component of spent media from *K. oxytoca* strains isolated from patients with AAHC was first reported in 1990 (29). With respect to *K. oxytoca* being a causative agent of AAHC, the bacterium has fulfilled Koch’s postulates (15). While a commensal of the gut microbiota of some individuals, it has been suggested that cytotoxic *K. oxytoca* is a transient member of the gut microbiota (29).

TV is a PBD produced by *K. oxytoca* and is a causative agent of AAHC (12). The TV biosynthesis genes are encoded on a non-ribosomal peptide synthase operon and include *npsA, thdA* and *npsB*. The genes *aroX* and *aroB* are also essential for TV production (13). *npsA*, *thdA, npsB* and *aroX* are located on a pathogenicity island (PAI). In clinical isolates, the PAI was present in 100 % of toxin-producing isolates, but only 13 % of non-toxin-producing isolates (12). AAHC is characterized by disruption of epithelial barrier function resulting from apoptosis of epithelial cells lining the colon. TV exerts its apoptotic effect by binding to tubulin and stabilising microtubules, leading to mitotic arrest (14).

A second PBD generated by the same pathway as TV has been identified (13). TM [also called kleboxymycin (11)] has stronger cytotoxic properties than TV, having a PBD motif with a hydroxyl group at the C11 position, while TV has an indole ring. When deprived of indole by the inactivation of the indole-producing tryptophanase gene *tnaA*, *K. oxytoca* produces TM but not TV. TV production is restored with the addition of indole, as indole spontaneously reacts with TM to produce TV. Limited interconversion between TM and TV may also occur spontaneously *in vivo* (11). TM is a genotoxin and triggers apoptosis by interacting with DNA, which leads to the activation of damage repair mechanisms, causing DNA strand breakage (14). DNA interaction is prevented in the case of TV by its indole ring, and both the molecular targets and apoptotic mechanisms of TM and TV are distinct. The kleboxymycin BGC is not native to *K. oxytoca*, nor the wider *Enterobacteriaceae*. Instead, the BGC is thought to have been acquired via horizontal gene transfer from *Xenorhabdus* spp., which in turn acquired the BGC from bacteria of the phylum *Actinobacteria* (11).

In the current study, we found the kleboxymycin BGC in our *K. michiganensis* isolates and that it was common among four species of the *K. oxytoca* complex, with *K. oxytoca* and *K. grimontii* strains making the largest contribution and the type strains of *K. grimontii* and *K. pasteurii* encoding the BGC (**Figure 3**). Prior to this study, sequences from two *K. oxytoca* strains (MH43-1, GenBank accession number MF401554 (11); AHC-6, GenBank accession number HG425356 (12)) were available for the kleboxymycin BGC. Draft genome sequences do not appear to be publicly available for either of these strains. However, our analysis of the kleboxymycin BGC across the *K. oxytoca* complex has shown that MH43-1 is a strain of *K. grimontii* (**Figure 3c**). Hubbard *et al.* (20) recently reported on a strain of *K. grimontii* that encoded the BGC, based on antiSMASH analysis. Comparison of the AHC-6 sequence with that of MH43-1 and other sequences included in this study shows AHC-6 is a strain of *K. oxytoca* (99.0–99.55 % nucleotide similarity with the BGCs encoded by the three *K. oxytoca* MAG sequences included in **Supplementary Figure 5**). It is likely that as more genomes of *K. oxytoca* complex species are deposited in public databases, the range of species encoding the kleboxymycin BGC will increase.

All three of our *K. michiganensis* strains encoded the kleboxymycin BGC (**Figures 2 and 3**), as did strains of *K. grimontii* we previously isolated from preterm infants and *K. oxytoca* and *K. michiganensis* MAGs recovered from publicly available shotgun metagenomic data (**Supplementary Figure 5**). Whether the BGCs encoded in our clinical and infant-associated strains are functional will be the subject of future studies. The discovery of the kleboxymycin BGC in strains and MAGs recovered from preterm infants is of particular concern. Gut colonization is linked with AAHC, with disease caused by the overgrowth of cytotoxin-producing strains secondary to use of antibiotics (16). AAHC presents as diffuse mucosal oedema and haemorrhagic erosions (16), and patients pass bloody diarrhoea (30). The gut microbiota of preterm infants is shaped by the large quantity of antibiotics these infants are given immediately after birth to cover possible early onset infection, with ‘blooms’ of bacteria preceding onset of infection (3). Blood in the stool is frequently associated with necrotizing enterocolitis (NEC) in preterm infants, which shares similar pathological hallmarks to AAHC – i.e. intestinal necrosis. Notably, NEC is difficult to diagnose in the early stages and is often associated with sudden serious deterioration in infant health, with treatment options limited due to emerging multi-drug-resistant bacteria associated with disease. Previous studies have linked *Klebsiella* spp. to preterm NEC (supported by corresponding clinical observations), with bacterial overgrowth in the intestine linked to pathological inflammatory cascades, facilitated by a ‘leaky’ epithelial barrier and LPS–TL4 activation. Recent work has demonstrated *K. oxytoca* complex isolates of ST173, ST246 and a novel ST (7-32-38-44-69-25-43) recovered from infants with NEC can produce kleboxymycin (TM) and TV (31). Using our MLST annotation scheme (**Supplementary Table 2**), we determine these sequence types represent *K. grimontii*, *K. grimontii* and *K. pasteurii*, respectively, with rMLST analyses of the whole-genome sequence data of Paveglio *et al.* (31) confirming our findings (rST 124484, rST 124487 and rST 157090, respectively). Taken together with the results from our study, we suggest specific virulence factors – i.e. kleboxymycin-related metabolites encoded by atypical *Klebsiella* spp. – may also play a role in NEC, and this warrants further study.

Attempts have been made to link specific subtypes of *K. oxytoca* to AAHC (32). Cytotoxic effects were limited to *K. oxytoca*, with faecal (and to a lesser extent skin) isolates of *K. oxytoca* most commonly associated with cytotoxicity (32). No genetic relationship was associated with cytotoxic strains based on pulsed-field gel electrophoresis, and 31/97 strains exhibited evidence of cytotoxin production (i.e. reduced viability of Hep2 cells). Joainig *et al.* (32) isolated genetically distinct cytotoxin-positive and -negative strains from one AAHC patient, leading them to suggest that, when detected in faeces, *K. oxytoca* should be considered an opportunistic pathogen able to produce disease upon antibiotic treatment. They also found that, in patients with acute or chronic diarrhoeal diseases, more than half of the isolates recovered were cytotoxin-positive. Given that *K. oxytoca*-related species are not routinely screened for in such samples, it is possible that kleboxymycin-producing isolates may make a greater contribution to diarrhoeal diseases than currently recognized, especially in patients suffering from non-*C. difficile*-associated disease. We have shown that there are species-specific differences in the kleboxymycin BGC (**Figure 3c**). These differences may have implications for virulence of strains and warrant further study. It is hoped that the identification of an increased range of strains (including type strains) encoding the kleboxymycin BGC will facilitate such studies.

## Abbreviations

AAHC: antibiotic-associated haemorrhagic colitis
AMR: antimicrobial resistance
BGC: biosynthetic gene cluster
MAG: metagenome-assembled genome
MLST: multi-locus sequencing typing
MSA: multiple-sequence alignment
NEC: necrotizing enterocolitis
PBD: pyrrolobenzodiazepine
rMLST: ribosomal MLST
TM: tilimycin
TV: tillivaline
VFDB: Virulence Factors of Pathogenic Bacteria Database

## ACKNOWLEDGEMENTS

Consultant microbiologist Dr Frances Davies (Imperial College Healthcare NHS Trust) is thanked for providing access to clinical strains. Imperial Health Charity is thanked for contributing to registration fees for the Professional Doctorate studies of PS. This work used the computing resources of CLIMB (33), the UK MEDical BIOinformatics partnership (UK Med-Bio, supported by Medical Research Council grant number MR/L01632X/1) and the Nottingham Trent University Hamilton High Performance Computing Cluster. The research placement of FM was funded by Nottingham Trent University. LJH is funded by a Wellcome Trust Investigator Award (no. 100/974/C/13/Z); a BBSRC Norwich Research Park Bioscience Doctoral Training grant no. BB/M011216/1 (supervisor LJH, student MK); an Institute Strategic Programme Gut Microbes and Health grant no. BB/R012490/1 and its constituent projects BBS/E/F/000PR10353 and BBS/E/F/000PR10356; and an Institute Strategic Programme Gut Health and Food Safety grant no. BB/J004529/1 to LJH. PS did all phenotypic characterization work. PS did initial bioinformatics analyses (assembly, annotation, CARD, initial BGC work), while FM and LH undertook the large-scale BGC analyses and infant-associated BGC work; LH undertook the MLST annotation work; ALM and MK prepared and pre-processed DNA for sequencing; LJH, ALM and LH supervised the work. We would like to thank Dave Baker and the QIB core sequencing team for WGS library preparation and sequencing. All authors contributed to interpretation and analyses of data, and writing of the manuscript.

